# Corticostriatal Plasticity Established by Initial Learning Persists After Behavioral Reversal

**DOI:** 10.1101/2020.04.16.045625

**Authors:** Sanchari Ghosh, Anthony M Zador

## Abstract

The neural mechanisms that allow animals to adapt their previously learned associations in response to changes in the environment remain poorly understood. To probe the synaptic mechanisms that mediate such adaptive behavior, we trained mice on an auditory-motor reversal task, and tracked changes in the strength of corticostriatal synapses associated with the formation of learned associations. Using a ChR2-based electrophysiological assay in acute striatal slices, we measured the strength of these synapses after animals learned to pair auditory stimuli with specific actions. Here we report that the pattern of synaptic strength initially established by learning remains unchanged even when the task contingencies are reversed. Our results suggest that synaptic changes associated with the initial acquisition of this task are not erased or over-written, and that behavioral reversal of learned associations may recruit a separate neural circuit.

## Introduction

One of the key neural mechanisms for adaptive behavior involves changes in the strengths of specific synaptic connections. Different behaviors involve different circuits, and thus recruit changes at different synaptic connections. In fear conditioning paradigms, for example, the association of a tone and a foot-shock induces freezing behavior that is mediated by long-term potentiation or LTP (Malinow and Malenka, 2002) at specific synapses that convey auditory information to the amygdala (LeDoux, 2000; Rumpel et al., 2005). Similarly, in barn owls the alignment of visual and auditory spatial maps for sound localization is mediated by specific synaptic connections in the inferior colliculus (Feldman and Knudsen, 1997). Although synaptic plasticity is thought to mediate many forms of learning, the specific loci of the synaptic changes have been experimentally established in only a handful of behavioral paradigms.

A hallmark of animal adaptation is that it is an ongoing and continual process, typically occurring not just once, but often throughout the lifetime of the animal. For example, a tone that predicts a shock one day might not predict it the next. It might seem intuitive that unlearning of such a previously formed association—”extinction”—would involve simply overwriting or erasing the synaptic changes underlying the initial tone-shock association. Indeed, optogenetic potentiation and depotentiation of auditory inputs to the amygdala can mediate bidirectional activation and deactivation of cue-induced freezing behavior (Nabavi et al., 2014). At the behavioral level, however, extinction of sound-induced freezing behavior does not appear to involve simple erasure of the initial memory, but rather inhibition of the freezing response by other brain structures (Quirk et al., 2000). Similarly, chronic prism placement alters the topography of synaptic connections in the inferior colliculus of the barn owl (Knudsen and Brainard, 1995) (Feldman and Knudsen, 1997), but the connections formed early in life persist even after they are no longer functionally expressed (Linkenhoker et al., 2005). By contrast, stimulation-induced persistent LTP in the hippocampus can be reversed if animals are exposed to novel environments as opposed to their familiar arena, suggesting that the same ensemble of synapses may be re-used in new environments (Xu et al., 1998). Thus, the extent to which ongoing behavioral adaptation to a changing environment recruits the same synapses as in the initial learning remains an open and complex question, the resolution of which may depend on the specific circuits and behaviors involved.

Many brain regions involved in learning are parts of the basal ganglia, which consist of distinct nuclei. The chief input nucleus—the striatum—integrates inputs from various cortical and sub-cortical areas. The motor striatum is broadly implicated in movement control (Klaus et al., 2019; Kreitzer and Malenka, 2008), and stimulating a particular subset of striatal neurons—the “direct pathway” neurons—in the anterior dorsal striatum (Kravitz et al., 2010) promotes contralateral movement. By contrast, the same stimulation in the auditory striatum (Guo et al., 2018), or in the dorso-medial striatum (Tai et al., 2012), only introduces a choice bias in the context of the behavioral task. Unlike the motor striatum, in which neuronal activity is finely tuned to movement initiation (Cui et al., 2013), neurons in the auditory striatal neurons mainly encode stimulus features during sound presentation (Guo et al., 2018). Moreover, recent studies show that dopaminergic projections to the posterior striatum originate from a different sub-part of substantia nigra compared to the anterior striatal regions and may not signal the commonly believed reward prediction error (Menegas et al., 2018; Menegas et al., 2015).

Here we have used an auditory two alternative choice decision task to study learning associated synaptic changes in a part of the posterior striatum—the auditory striatum. Training rats to perform an auditory discrimination task (Znamenskiy and Zador, 2013) induces potentiation of corticostriatal synapses, forming a spatial plasticity gradient along the tonotopic gradient of auditory inputs to the auditory striatum (Xiong et al., 2015). The sign of this gradient, which can be read out in acute slices of the auditory striatum, is determined by the precise stimulus-response association learned: in animals trained to associate low frequency stimuli with a left decision, the sign of the gradient is opposite to that in animals trained to associate low frequency stimuli with a right decision (Xiong et al., 2015). These and other observations suggest that strengthening connections between sensory cortices and their striatal targets facilitates appropriate action selection after learning (Guo et al., 2018; Kreitzer and Malenka, 2008; Lee et al., 2015). In this study, we exploit our high-resolution understanding of the synaptic changes elicited by acquisition of this behavior to test whether reversal of the stimulus-response contingencies leads to reversal of the corresponding synaptic strengths. We find that the plasticity gradient established by the initial training in this auditory task is not modulated bidirectionally, but instead remains stable even after the contingencies are reversed.

## Results

To assess the synaptic changes associated with acquisition and subsequent reversal of stimulus-response contingencies, we tested the effect of reversal learning on the plasticity gradient of corticostriatal projections in mice. In rats, this gradient is such a sensitive measure of learning contingencies that it can reveal, with 100% accuracy, whether an individual subject has been trained to associate a high-frequency stimulus with a left or a right choice (Xiong et al., 2015). Because the present experiments were conducted in mice, we first confirmed that they can also be rapidly and reliably trained to perform the two alternative choice (2-AC) tonecloud task (Chen et al., 2019); see also (Guo et al., 2018; Jaramillo and Zador, 2014). We then tested whether they could be reliably trained to reverse the contingencies upon which they had initially been trained. Next, we assessed whether projections from mouse cortical area A1 to striatum are organized in a tonotopic fashion. Taking advantage of this tonotopic organization of the corticostriatal projection, we used an electrophysiological assay to assess strength of these synapses along the tonotopic axis. This ability to read out the synaptic correlate of the learned association provides a unique opportunity to probe the changes in this circuit after behavioral reversal. Finally, using the reversal paradigm, we show that the plasticity gradient established at auditory corticostriatal synapses after the initial learning phase persists even after successful reversal learning. Our results suggest that the site of plasticity engaged during reversal may differ from that engaged in the initial learning of the task.

### Acquisition time for initial association and reversal are comparable

The auditory 2-AC task, adapted from a related task developed for rats (Znamenskiy and Zador, 2013), required that subjects discriminate between low and high “tonecloud” stimuli, and report their choice by going to either the left or right choice port (Fig. 1A). Subjects initiated a trial after the ‘Go’ cue (light ‘on’ at center port) was provided. On each trial, the stimulus consisted of a train of short overlapping pure tones drawn from either a low (5-10kHz) or high (20-40kHz) octave. Subjects were required to listen to the entire stimulus (500ms) before reporting their choice; at the end of the stimulus they were rewarded with a small drop of water (0.5ul) at the center port, to encourage them to remain in the center port for the duration of the stimulus. After withdrawal from the center port, subjects were required to choose between a left and a right reward port, depending on the frequency content of the stimulus (Fig. 1A, top). Mice readily learned this task over a period of 2-3 weeks.

**Figure 1.**
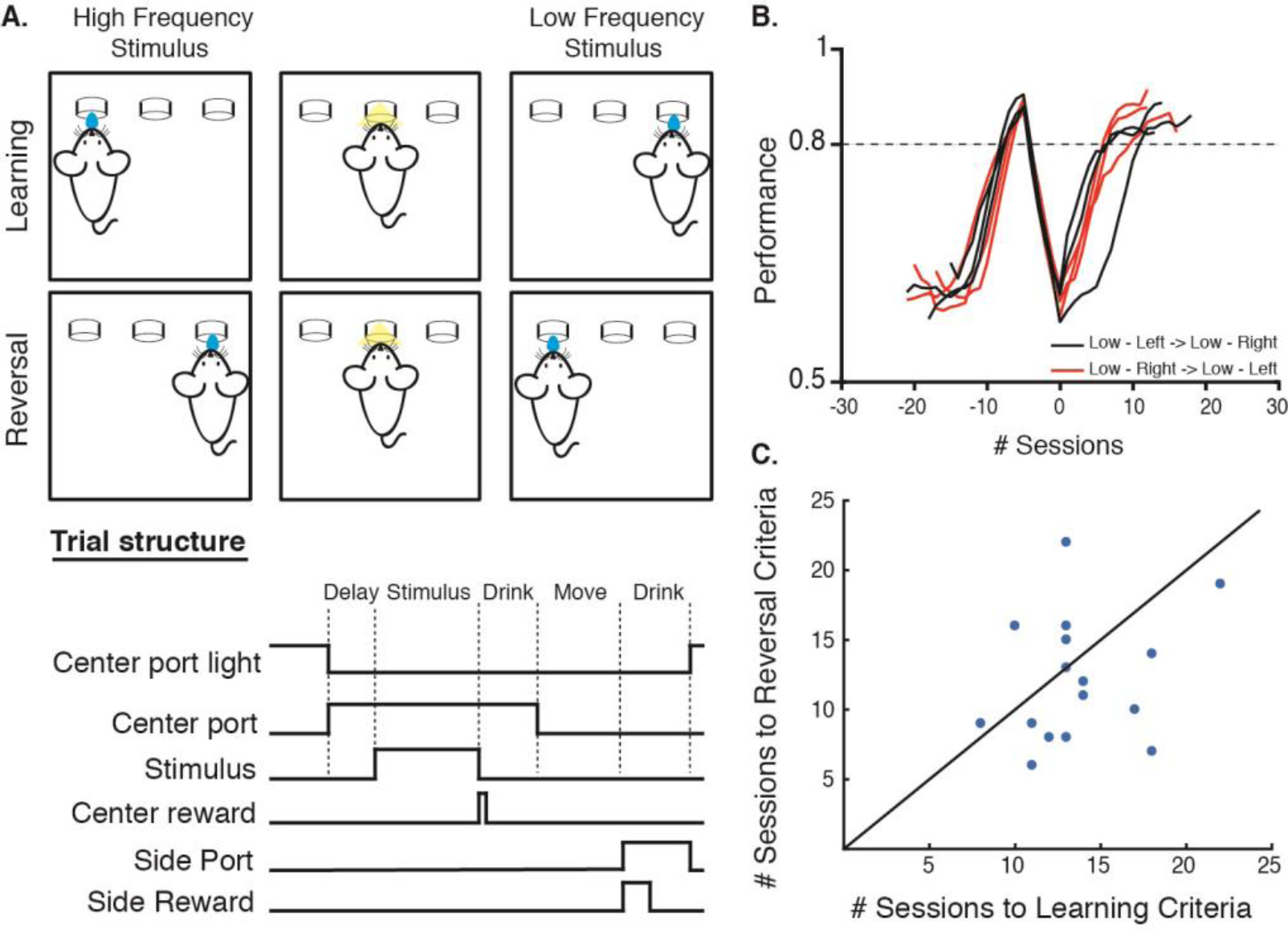
Training mice on a reversal paradigm based on 2-AC frequency discrimination. **(A)** The schematic of the reversal paradigm using the tonecloud task for mice (*top*). Animals are first trained on one contingency, e.g., to pair a Low frequency tonecloud with reward on the right referred to as ‘Low-Right (‘Learning’). Once they reach the performance criteria, the training contingency is reversed, requiring the same animal to now pair a Low frequency tonecloud with reward on the left, or, Low-Left (Reversal). The trial structure (*bottom*) shows the sequence of events in a single typical ‘correct’ trial in the task. **(B)** Example learning curves of mice trained in the reversal paradigm (*black*: Low-Left → Low-Right, n=3 and *red*: Low-Right → Low-Left, n=3), where 0 denotes the start of training on the reversed contingency. The performance of the animals is smoothed over 3 sessions for this plot. **(C)** Animals require a comparable number of sessions to reach performance criteria during initial ‘Learning’ as during ‘Reversal’.

We next established a reversal paradigm in which subjects trained to criterion on one association (Low-Left or Low-Right) were then trained to reverse this association (Fig. 1A, bottom). To avoid overtraining subjects on one contingency, and thereby potentially increasing the difficulty in re-training them on the reversed contingency, we established a relatively lax performance criterion of >80% correct per session. After 4-6 sessions of >80% performance, we reversed the stimulus-response contingency. In the sessions immediately following reversal, all subjects show a marked decrease in performance, often performing well below chance, after which performance increased to levels comparable to the original contingency (Fig.1B). On average, subjects required a similar number of sessions to reach the fixed performance criteria (12.9 ± 0.8 (SEM), vs. 12.5 ± 1.2 (SEM), p=0.61, Wilcoxon Signed Rank test) (Fig.1C). Thus, mice could be reliably trained to perform both the basic tonecloud task and the reversal.

### Auditory corticostriatal projections in mice show tonotopy

The mouse auditory system is organized tonotopically (Hackett et al., 2011; Kaas, 2010; Kanold et al., 2014). This feature allows measuring neural activity and plasticity in specific tone-responsive regions (Bathellier et al., 2012; Higgins et al., 2010; Kaja Ewa Moczulska, 2013). We mapped the tonotopic projections from primary auditory cortex to auditory striatum. We first performed intrinsic optical imaging of the auditory cortex through a thinned bone preparation in a mouse (Bakin et al., 1996; Bathellier et al., 2012), in response to pure tones of 4kHz and 32kHz (Fig. 2A). The intrinsic signals elicited by these stimuli consistently revealed three regions, which we identified as A1, A2 and AAF (Fig. 2B), and subsequently mapped to the brain surface using the vasculature as guidance. We then performed small focal injections in high and low frequency regions of A1, using AAV1-CAG-GFP and AAV1-CAG-tdTomato respectively (Fig. 2A, supplementary Fig. S1). Inspection of coronal sections of the auditory striatum revealed a tonotopic organization of the afferent cortical projections (Fig. 2C). Fibers from the low frequency region of A1 terminated more medially, whereas those from the high frequency region terminated more laterally (Fig. 2D & E). These experiments reveal a tonotopic projection in the mouse from A1 to the auditory striatum, which can be observed in standard coronal slices.

**Figure 2.**
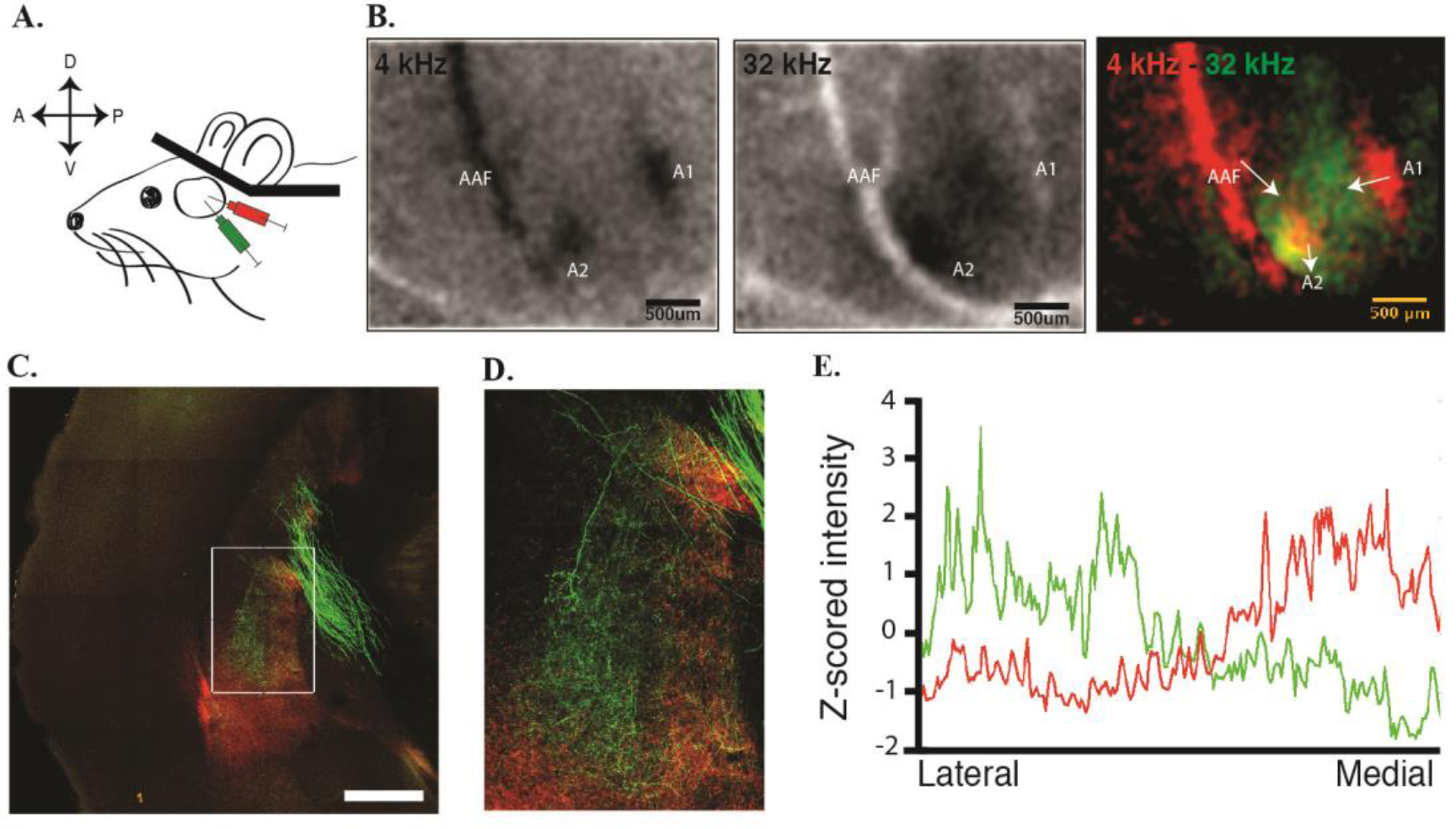
Auditory corticostriatal projections in mice are tonotopically organized. (**A**) Intrinsic optical imaging of the auditory cortex in a head-fixed mouse through a window of thinned bone. (**B**) Intrinsic optical images in response to pure tones of 4 and 32kHz (left and middle). Composite image showing tonotopy in auditory cortex where green corresponds to high frequency and red to low frequency. The arrows indicate overall tonotopic gradient (low to high) in the individual tone responsive areas. (**C**) Tonotopic separation of A1 projections into auditory striatum showing high frequency A1 projects more laterally and low frequency A1 projects more medially (white rectangle). Scale bar=500um. (**D**) Enlarged view of auditory striatum (white rectangle) showing tonotopic separation quantified in **E**.

### Training induces a tonotopic gradient of synaptic strength in the auditory striatum

Striatal plasticity has been previously implicated in skill learning and associative learning (Cox and Witten, 2019; Pasupathy and Miller, 2005; Yin et al., 2005). In rats, acquisition of a frequency-dependent auditory task establishes a gradient of synaptic strength along the tonotopic gradient in striatum (Xiong et al., 2015). We tested whether learning the tonecloud task also resulted in a stereotyped gradient of corticostriatal synaptic strength in the mouse striatum.

We used the channelrhodopsin-2-evoked local field potential (ChR2-LFP) in acute slices of auditory striatum to measure the strength of the corticostriatal synapses at specific locations along the striatal tonotopic gradient (Xiong et al., 2015). To ensure that the ChR2-LFP selectively reflected the strength of cortical, rather than thalamic or other inputs to the striatum, we expressed AAV9-CAG-ChR2 in the primary auditory cortex (Fig. 3A). Animals were approximately 5 weeks old at the time of injection. After 3-5 days of recovery, the animals were trained on the tonecloud task for about 2-3 weeks until they reached the performance criterion. The duration of the training also allowed for the expression of ChR2 in the infected neurons. Once an animal reached the behavioral performance criterion (above 80% for 4-6 consecutive sessions), we obtained acute coronal slices (Fig 3A, top right) from its brain, and recorded the ChR2-LFP from the auditory striatum (Fig. 3B).

**Figure 3.**
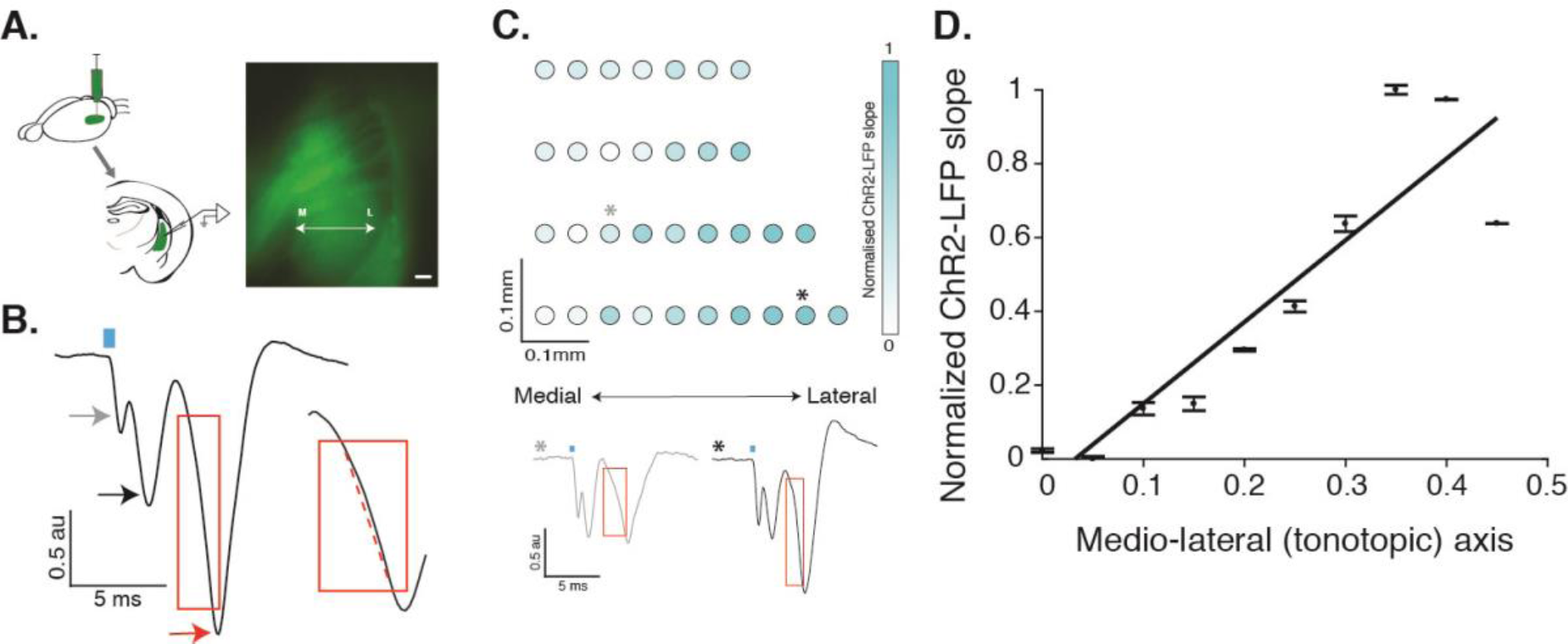
ChR2-LFP slope measurements reflect the learning induced plasticity gradient. (**A)** AAV9-ChR2 is injected in auditory cortex and recordings are obtained from acute coronal slices of auditory striatum exhibiting ChR2-expressing corticostriatal fiber terminals. (*Right)* Example of an acute slice showing ChR2-GFP expressed in corticostriatal fibers. (**B**) Example trace of ChR2-LFP from one position in the slice. *Gray arrow* indicates a light artefact often observed soon after laser stimulation (*blue rectangle*). *Black arrow* indicates the depolarization of ChR2-expressing corticostriatal fibers. *Red arrow* indicates the post-synaptic response of downstream striatal neurons. The response is normalized to the fiber depolarization, and the normalized ChR2-LFP slope is calculated from the post-synaptic component (*red rectangle*). Inset shows calculation of ChR2-LFP slope by fitting a line (*dotted red line*) to the post-synaptic component. (**C**) Representative image showing distribution of individual normalized ChR2-LFP slopes along the tonotopic axis of the left auditory striatum of an example animal trained on Low-Left contingency. Two example traces corresponding to two data points on top are shown in the bottom of panel C. The *red rectangle* encloses the initial depolarization phase showing a faster depolarization for the lateral data point (*black \**) compared to the medial one (*grey \**). (**D**) Mean and standard deviation of the ChR2-LFP slope data from C, plotted along the tonotopic axis. Slope of the linear fit to these data points is the plasticity gradient (= 0.33) for this animal.

ChR2-LFPs evoked in these slices showed a stereotyped waveform, reminiscent of that seen in classic extracellular LFPs evoked by electrical stimulation of the Schaeffer collateral input to the CA1 region of the hippocampus (Xiong et al, 2015). Because the striatum, like the hippocampus, lacks recurrent excitatory connections (Kreitzer and Malenka, 2008), this LFP can be used as a measure of synaptic efficacy (Xiong et al, 2015). As expected, the ChR2-LFP responses were evoked only in regions containing ChR2-expressing fibers (supplementary Fig. S2A), and the magnitude of the ChR2-LFP increased with duration and strength of optical stimulation (supplementary Fig. S2B & C). Pharmacological dissection of the stereotyped ChR2-LFP waveform uncovered three distinct components. The first was a very short latency light artefact (arising from the photoelectric effect); the second was the fiber volley; and the third was the synaptic response, sensitive to blockage by the AMPA receptor antagonist CNQX (supplementary Fig. S2D). The slope of this third CNQX-sensitive component—the ChR2-LFP slope— represented a measure of corticostriatal input (Fig. 3B).

We recorded the ChR2-LFP from multiple positions within the left auditory striatum in each slice, and calculated the ChR2-LFP slope at each position (Fig. 3C, top). The slope of these ChR2-LFP values along the tonotopic axis represents the plasticity gradient for each animal (Fig 3D). Based on the anatomical projections of the tonotopic inputs from the cortex to the striatum, we expected a larger ChR2-LFP in the lateral auditory striatum for animals trained on the Low-Left contingency, and lower for animals trained on the opposite (Low-Right) contingency. Fig. 3D shows the gradient along the medio-lateral (tonotopic) axis of a single animal trained to associate low frequency sounds with left decisions (Low-Left). As expected, the ChR2-LFP slope is positive (p = 0.0004, for positive correlation along mediolateral axis). This result was reliable: All animals (8/8) trained on the Low-Left contingency showed a positive slope (Fig. 4B, cyan bar; mean plasticity gradient = 0.19 ± 0.03 (SEM), each circle represents individual animals). By contrast, all (8/8) animals trained on the opposite (Low-Right) contingency showed a negative slope (Fig. 4B, cyan bar; mean plasticity gradient = −0.19 ± 0.04 (SEM), each square represents individual animals). By contrast, there was no gradient along the dorsoventral axis of the striatum, consistent with the fact that inputs along this axis are not organized tonotopically (supplementary Fig. S3). Thus, training mice to associate a low or high frequency sound with a right or left choice reliably establishes a robust gradient of synaptic strength along the tonotopic axis that faithfully represents the learned sound-action contingency (p=8 × 10^−5^ between plasticity gradients of animals trained on Low-Left vs Low-Right, Wilcoxon Rank Sum test, n=8 in each group).

**Figure 4.**
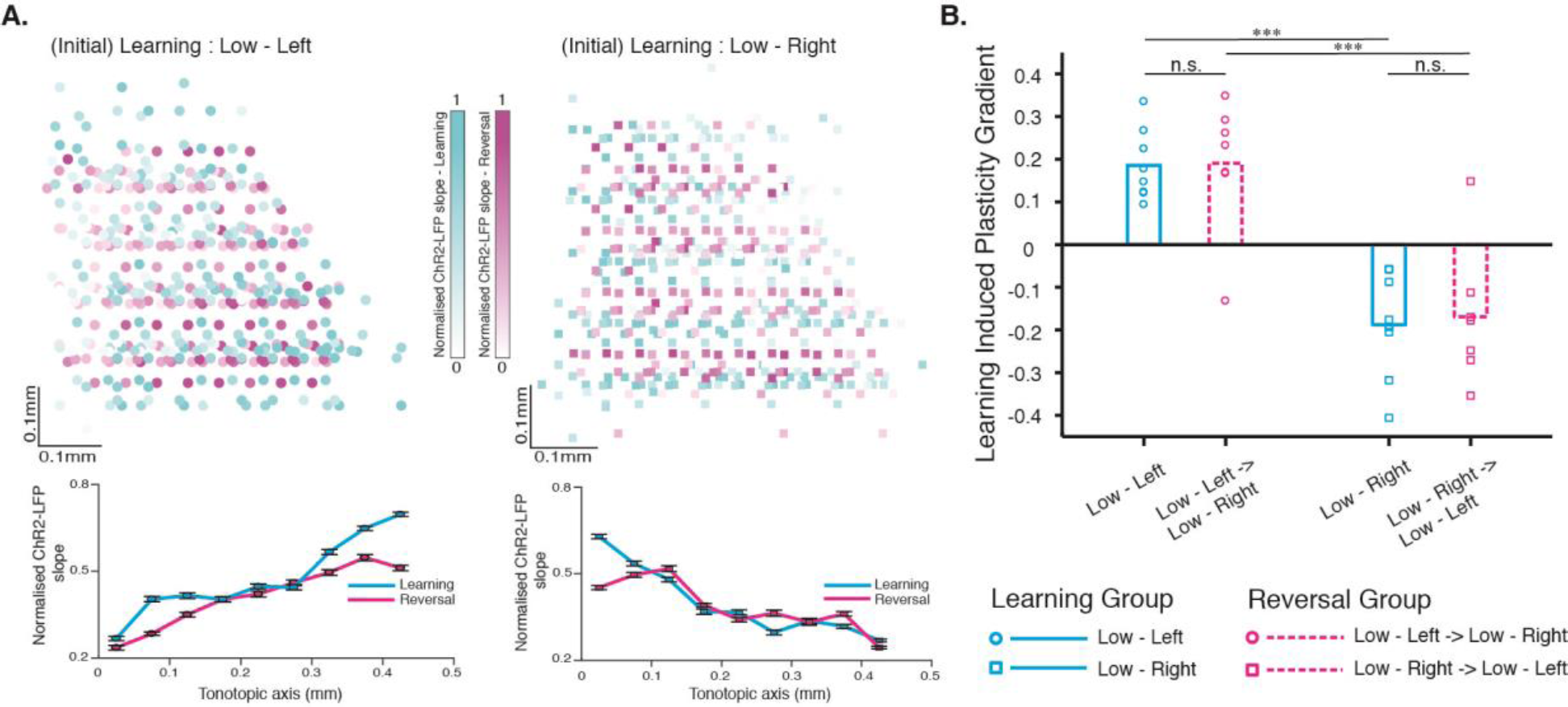
Learning-induced plasticity gradient in auditory corticostriatal circuit reflects initial learning contingency. (**A**) *Top* - ChR2-LFP slope after either learning alone (*cyan*, n=8 animals) or reversal (*magenta*, n=7 animals) from Low-Left (*left*) or Low-Right (*right*) contingency on the tonecloud task. The intensity of each point represents magnitude of normalized ChR2-LFP slope value recorded at that position of the striatal slice. *Bottom* - mean ± SEM vs. tonotopy. (**B**). Summary of learning-induced plasticity gradient from 4 groups of animals: {Learning (*cyan*) or Reversal (*magenta*)} x {Low-Left (*circle*) or Low-Right(*square*)}. Points represent individual animals. Significant differences were only observed between groups trained on opposite contingencies: Low-Left vs. Low-Right (p=8 × 10-5) and Low-Left→Low-Right vs. Low-Right→Low-Left (p=2 × 10-3); Wilcoxon Rank Sum test.

### The plasticity gradient established during initial training persists even after animals are trained to reverse the learned sensory-motor association

The comparable training times required by the animals to reach criterion performance during learning and reversal (Fig. 1B) would suggest significant changes in synaptic strengths in the neural circuit. In our simple working model, the strengthening of specific synapses from the auditory cortex to the auditory striatum is one of the circuit changes underlying the acquisition of the tonecloud task. In particular, the fact that these synapses are strengthened along the striatal tonotopic gradient, and that the sign of the gradient depends on the specific contingencies (Low-Left or Low-Right) acquired during learning (Fig. 4), suggested a simple prediction: reversing the contingencies should reverse the sign of the gradient of corticostriatal synaptic strength along the tonotopic axis.

We therefore tested the effect of reversal on corticostriatal plasticity. For each animal injected with ChR2 and trained to criterion on one contingency, we reversed the contingencies and re-trained to criterion. To minimize bias, we performed recordings (blind to the training contingency).

Surprisingly, contrary to the predictions of the simple model, we found that reversing the association did not alter the sign of the plasticity gradient. The sign of the gradient was negative in animals initially trained on the Low-Right contingency, and remained negative even if they were subsequently trained to Low-Left (Fig. 4A, right, magenta). Similarly, sign of the gradient was positive in animals initially trained on the Low-Left contingency, even if they were subsequently trained to Low-Right (Fig. 4A, left, magenta). These results were robust and consistent across animals: In 100% (16/16) of animals trained without reversal, and 86% (12/14) animals trained on reversal, recordings from a single brain slice could be used to infer the animal’s initial training history. Thus, our results demonstrate that the plasticity gradient established during the initial learning persists even after subsequent learning of the opposite association.

## Discussion

We have investigated how the auditory corticostriatal circuit, critical for auditory discrimination behavior, adapts to learning and reversal of stimulus-action association. Our main findings are that (1) learning in this task establishes a strong plasticity gradient in these synapses, in a pattern that reflects the training contingencies (Fig. 3E; (Xiong et al., 2015)); and (2) training animals to reverse this association leaves this initial gradient intact. Our observations have implications for the role of corticostriatal plasticity in mediating stimulus-action associations, and more broadly, for understanding how animals adapt to an ever-changing world.

The auditory striatum, located at the caudal tip of the striatum in the rodent, receives convergent input from auditory cortex, auditory thalamus and midbrain dopamine neurons. In animals performing an auditory task, inactivation of either auditory cortical or thalamic inputs to the auditory striatum (Chen et al., 2019), or of the auditory striatum itself (Guo et al., 2018) markedly impairs performance. Optogenetic activation of cortical inputs to the striatum elicits a choice bias that depends on the frequency tuning of the stimulated site (Znamenskiy and Zador, 2013). Acquisition of the tonecloud discrimination task strengthens corticostriatal inputs during learning, in a frequency specific manner (Xiong et al., 2015). Taken together, these results suggested a simple model in which the auditory striatum couples sensory inputs to rewarded actions, in which this coupling is mediated by the specific pattern of synaptic strength of cortical inputs to the striatum (Fig. 3; (Xiong et al., 2015)).

The present results argue against this simple model. If the auditory striatum were simply transforming auditory sensory information from the cortex into an action, then reversing the stimulus-action association would be predicted to reverse the gradient of synaptic strength in the striatum. However, we did not observe such a reversal in the plasticity gradient. Instead, we found that the gradient in synaptic strength depended only on the initial contingencies of the task the animal was trained to perform. In principle, the thalamic inputs to auditory striatum might play a role in transforming auditory sensory information into action, although the fact that the major thalamic inputs to the striatum arise from the dorsal medial geniculate nucleus (Chen et al., 2019)—an area in which neurons are not well-tuned to sound—argues against these inputs providing the necessary frequency-specific information. Furthermore, electrophysiological recordings in behaving mice show that the activity of auditory striatal neurons was only modestly influenced by the animals’ choices (Guo et al., 2018). Taken together, these observations suggest that the locus for transforming sensory inputs to actions is likely outside of the auditory striatum.

Our results indicate that rather than over-writing the initial memory trace, animals may keep this initial trace intact. This strategy may prevent what in artificial neural network research has been termed “catastrophic forgetting” (Michael McCloskey, 1989): the loss of old memories upon acquisition of new ones. In some cases, such as the alignment of auditory and visual maps, it appears that several distinct alignments can co-exist within a single circuit (Feldman and Knudsen, 1997; Knudsen and Brainard, 1995; Linkenhoker et al., 2005). However, a more general solution to the catastrophic forgetting problem may be to recruit new brain regions when a new memory is added. In the case of the reversal of stimulus-reward associations (Schoenbaum et al., 2009), several brain areas have been implicated, including the orbitofrontal cortex, the medial prefrontal cortex, the basolateral amygdala and the ventral striatum (Johnson et al., 2016; Schoenbaum et al., 1999; Schoenbaum and Setlow, 2003). Unraveling the circuit and synaptic basis of reversal learning in this task may provide a foundation for understanding how both natural and artificial systems adapt to new situations whilst avoiding catastrophic forgetting.

## Author Contributions

S.G. and A.M.Z. designed the study. S.G. performed experiments and analyzed data. S.G. and A.M.Z. wrote the manuscript.

## Acknowledgements

The authors acknowledge NIH and the Charles A. Dana Fellowship from The Dana Foundation, for funding this research. We would like to thank Federico Carnevale, Fred Marbach, Barry Burbach, Ashlan Reid and Hysell Oviedo for technical assistance. We also acknowledge constructive inputs from Arkarup Banerjee, Anqi Zhang, Christopher Krasniak, Dennis Maharjan, and other members of the Zador lab while preparing this manuscript.

## Supplementary Information

**Supplementary Figure S1.**
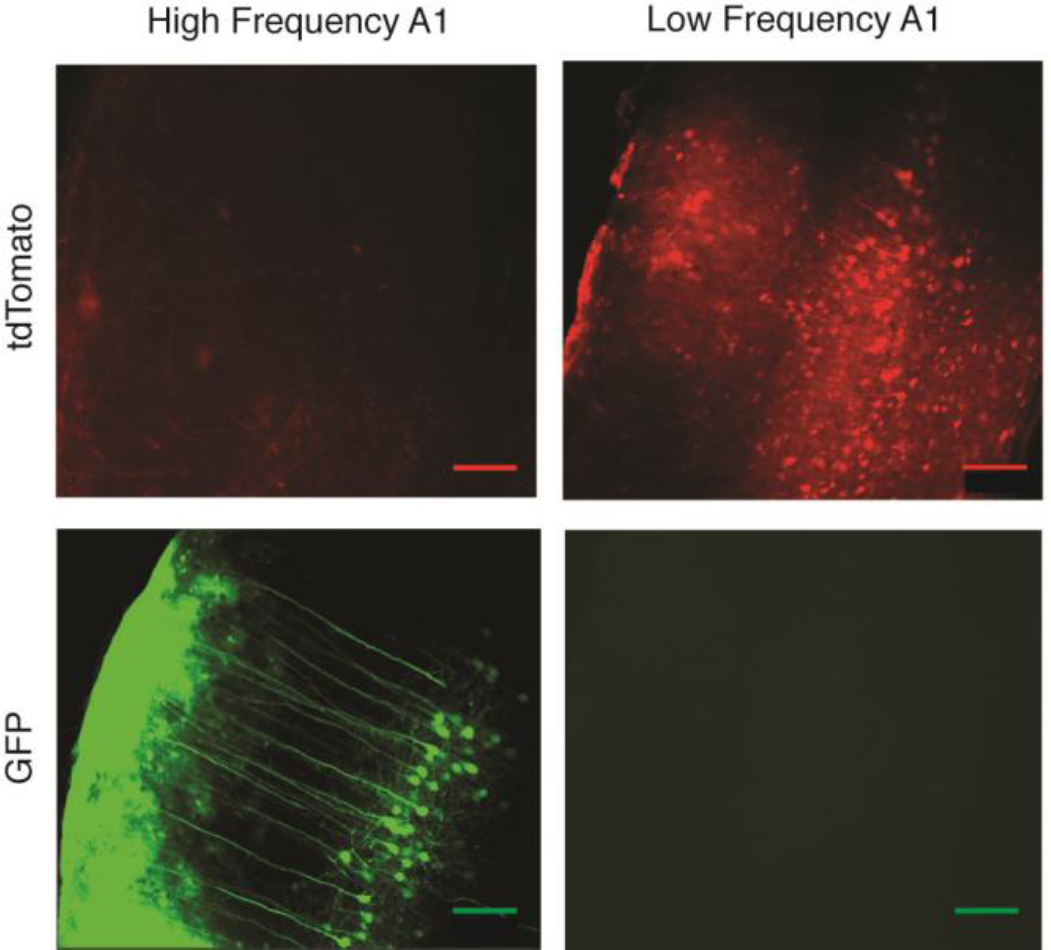
Injection site in primary auditory cortex confirming expression of tdTomato (*top*) and GFP (*bottom*). These images confirm little to no overlap of viral infections at the cortical injection site. Scale bar = 500μm.

**Supplementary Figure S2.**
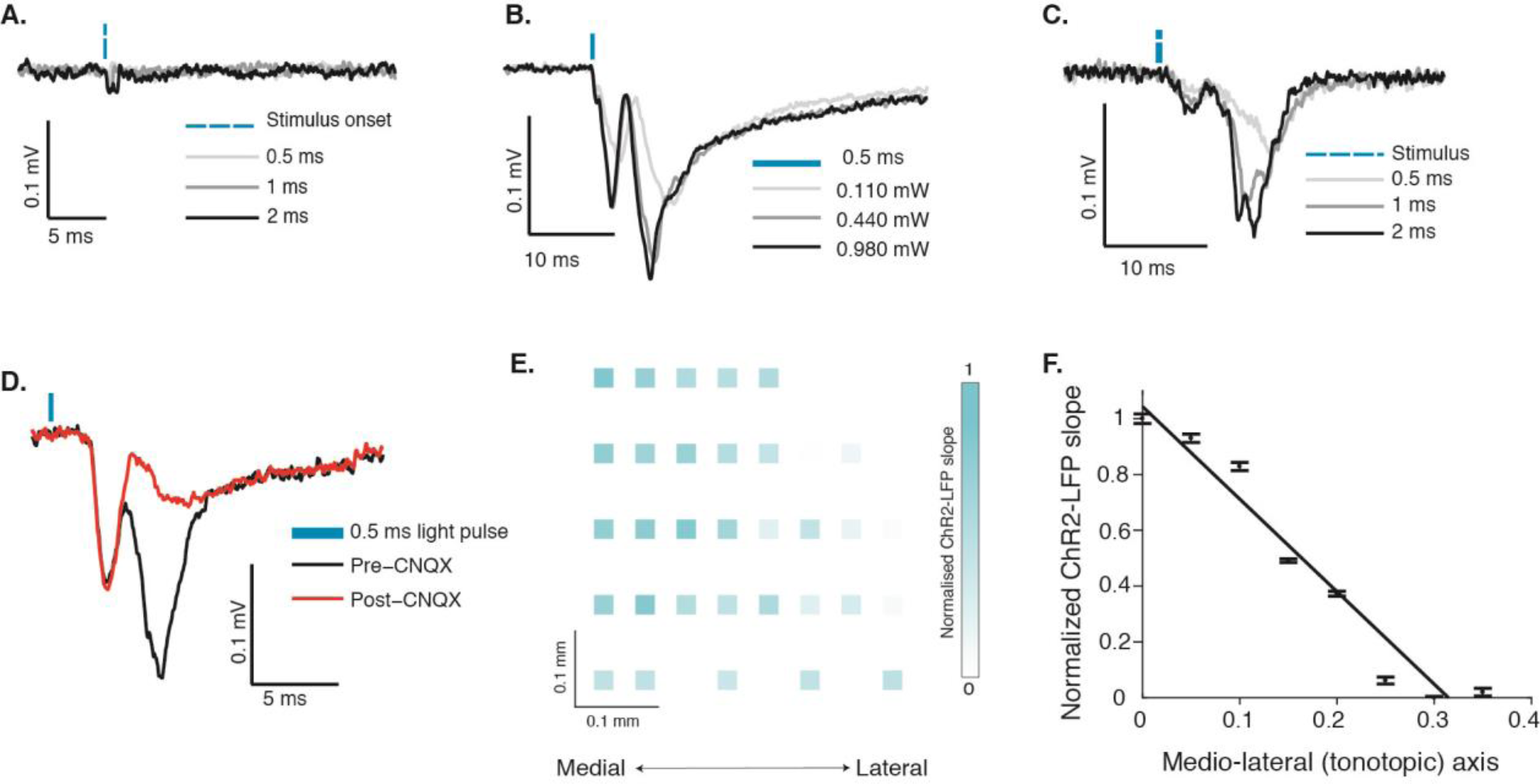
Controls of ChR2-LFP recordings and measurement of ChR2-LFP slopes. (**A**). Neuronal responses to optical stimulation is absent in brain region not expressing ChR2 (somatosensory cortex). (**B**). The magnitude of ChR2-LFP increases with increase in laser power, keeping the duration of stimulation at 0.5ms. (**C**). The magnitude of ChR2-LFP increases if duration of stimulation is increased at the highest laser power of 0.980mW. D. 30 mins of slice incubation with 50μM CNQX abolishes the post-synaptic response of striatal neurons without affecting the depolarization of cortical fiber terminals in striatum (*red*) in comparison to pre-drug control (*black*). (**E**). Example of normalized ChR2-LFP slope distribution in the left auditory striatum of an animal trained on the Low-Right contingency. (**F**). Mean and SD of the normalized ChR2-slope data from (**E**) plotted along the tonotopic axis.

**Supplementary Figure S3.**
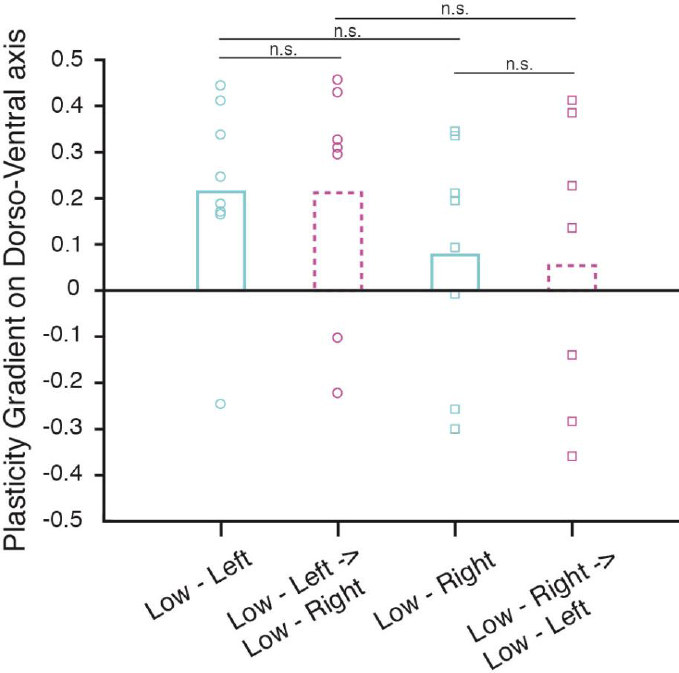
Summary of normalized plasticity gradient calculated along the dorso-ventral axis (non-tonotopic axis) does not reflect a consistent difference between training contingencies (Low-Left vs. Low-Right) or across training phases (learning vs. reversal). Kruskal

## Methods

### Animals and Viruses

All procedures were conducted in accordance with the institutional animal use and care policies of CSHL and NIH. C57 Black6J animals were obtained from Jackson laboratories and housed in a temperature and moisture controlled room with 12-hour light/dark cycle. Viruses used for anatomical tracing experiments – AAV2.1.CAG.GFP and AAV2.1.CAG.tdTomato were ordered from University of Pennsylvania Vector Core. AAV2.9.CAG.Channelrhodopsin virus for optogenetic stimulation was obtained from UNC Vector core.

### Surgical procedures and Injections

For performing stereotaxic injections, mice were anaesthetized with a cocktail of ketamine (60mg/kg) and medetomidine (0.5mg/kg) and immobilized on a stereotaxic set up. After sterilization with 70% alcohol and numbing with subcutaneous injection of Lidocaine (2mg/kg), the skin and tissue overlying the left auditory cortex was dissected to expose the temporal bone. To cover the entire primary auditory cortex, two injections were made perpendicularly to the brain surface roughly 2 mm and 2.5 mm caudal to the temporoparietal suture, and roughly 1 mm below the ventral edge. Each injection was made at two depths (400 μm and 600 μm) releasing approximately 80nl of virus at each depth.

### Transcranial intrinsic optical imaging

Mice were anesthetized with ketamine (60 mg/kg) and medetomidine (0.5 mg/kg) and immobilized in a stereotaxic setup. After sterilization with 70% alcohol and numbing with subcutaneous injection of Lidocaine (2 mg/kg), a portion of the scalp on the top of the cranium was removed and a headbar was attached to the exposed skull using Metabond adhesive (Parkell, S380), further secured using dental cement (Lang, Jet denture repair powder/liquid). After 2-3 days of recovery, the animals were anesthetized using Isofluorane (2.5% Isofluorane + Oxygen at 0.1 L/min) and immobilized in the stereotaxic set up using the headbar. After sterilization with 70% alcohol and numbing with subcutaneous injection of Lidocaine (2 mg/kg), the skin and tissue overlying the left auditory cortex was removed exposing the bone surface. For transcranial imaging, the bone was thinned using a low-speed dental drill. The exposed skull was kept moist during the imaging session using 1.5% agar in PBS. Additional anesthesia was provided by injecting Chlorprothixene (0.7mg/kg) and the mouse was then transferred for intrinsic optical imaging to a custom-built microscope set up. During the imaging, the mouse was kept lightly anaesthetized using isofluorane (1%Isofluorane + Oxygen at 0.1 L/min) while placed on a temperature regulated heating pad to maintain the body temperature close to 35°C. Intrinsic signal images were acquired using a CCD camera (Vosskuehler 1300QF) after illumination using red LED (615 nm). In order to evoke stimulus responses in the auditory cortex, 1s pure tone pips of were played at an interval of 30s. The frequencies chosen were 4kHz and 32kHz, each being presented at least 15 times at 80db. The acquired images were analyzed to depict normalized difference of reflectance in response to the stimulus ((pre-stimulus – post-stimulus)/pre-stimulus). The location of specific tone-responsive regions in these images were registered with respect to the image of the surface vasculature acquired using blue LED (488nm). These maps were subsequently used to perform tonotopic tracing experiments.

### Behavioral training and apparatus

Water-deprived animals were trained in custom sound-booths by Industrial Acoustics Company (Bronx, New York) containing a custom-built behavioral arena (20cm x 20 cm x 20 cm). This consisted of 3 ports located on one wall with inter port distance of 5.5cm (center to center). The height of the port was 2.5cm from the floor. The side walls of the arena had perforations aligned to the speakers located just outside the walls. Water was delivered through the ports via 19 gauge stainless steel tubes connected to rubber tubing (Silastic) and the flow was controlled via solenoid valves (Lee company). The valve opening times were calibrated at regular intervals to ensure accurate delivery of 0.5μl and 2.5μl of water from the center and side ports, respectively. LEDs located just above the water ports were used to provide ‘Go’ cues. The auditory stimuli were designed by the MATLAB protocol and delivered through the speakers that were calibrated described before (Jaramillo and Zador, 2014). The behavior system was automated and controlled through custom software written in MATLAB to operate the state machine interface of the behavior control module – bpod (https://sanworks.io/shop/viewproduct?productID=1027). In stage 1 of training, animals poked at the center port in response to a steady center port light-the ‘Go’ cue for trial initiation. Holding at center port for 50ms of pre-stimulus delay successfully initiated a trial and triggered delivery of the sound stimulus and a small reward (0.5ul) at the center port. A steady light at the correct side port (depending on the frequency content of the sound stimulus) signaled the mouse where to go next for an additional 2.5ul water reward within the trial duration of 10s. A new trial started when the animal reported its choice or if the trial time elapsed. There were no punishments for incorrect choices or early withdrawals at this stage. The animals were promoted to stage 2 when they completed more than 100 successfully rewarded trials within one session in stage 1. In stage 2, both the side port lights were turned on after stimulus delivery to signal the animal that it was time to report a choice and no longer signaled the correct port. At this stage, animals were given a white noise punishment for incorrect trials and early withdrawals. The animals moved on to stage 3 after 2 sessions of more than 100 completed trials each. The stage 3 is where animals spent maximum time in training. At this stage, the center port light still provided a ‘Go’ cue for trial initiation but side port lights stayed off. Early withdrawals were punished with a 1s time out and incorrect choices with a 4s time-out in addition to the white noise. At this stage the pre-stimulus delay was also increased to 250ms. Animals were either trained to pair low frequency toneclouds with a leftward movement and high frequency toneclouds with rightward (referred to as Low-Left) or vice versa (referred to as Low-Right). The training contingency for each animal (Low-Right or Low-Left) was randomly pre-determined by the experimenter. Animals were trained to the performance criteria of higher than 80% in 4-6 consecutive sessions before proceeding to recording experiments. For animals performing the reversal task, reversal of contingency was introduced after the same performance criteria as above while maintaining task parameters of stage 3. These animals were then subsequently trained to the same performance criteria in the opposite contingency before proceeding to recording experiments.

### Slice Experiments

Mice were first anaesthetized with a cocktail of ketamine (60mg/kg) and medetomidine (0.5mg/kg) then perfused with ice-cold artificial cerebrospinal fluid (aCSF) bubbled with 95% oxygen and 5% CO_2_. The mouse was then rapidly decapitated and the brain was removed from the cranium and placed in ice-cold cutting buffer also bubbled with 95% oxygen and 5% CO_2_. It was then transferred to the stage of a vibratome kept submerged in ice-cold cutting buffer (110mM choline chloride, 25mM NaHCO3, 25mM D-glucose, 11.6mM sodium ascorbate, 7mM MgCl2, 3.1mM sodium pyruvate, 2.5mM KCl, 1.25mM NaH2PO4, and 0.5mM CaCl2) continuously bubbling with 95% oxygen and 5% CO_2_. The temperature of the entire set up was maintained at 4°C. The brain was then cut into coronal slices of 250μm thickness until we reached the canonical slice chosen for our striatal recording. Once the ideal slice was cut, it was quickly transferred into a holding chamber containing continuously aerated aCSF (127mM NaCl, 25mM NaHCO3, 25mM D-glucose, 2.5mM KCl, 4mM MgCl2, 1mM CaCl2, and 1.25mM NaH2PO4, aerated with 95% O2 & 5% CO2) at 32°C. The slice was allowed to recover for about 30 mins and then maintained at room temperature at which the recordings were performed. After recovery, the slice was carefully transferred to the recording set up. CNQX was added to a final concentration of 50μM in aCSF and delivered through the perfusion system for inactivating glutamatergic synaptic transmission at corticostriatal synapses.

### Electrophysiology recordings and analysis

Local Field Potentials (LFPs) were recorded using Axopatch 200B amplifiers (Axons Instruments, Molecular Devices) using thin walled glass pipettes of resistance 2-3MΩ filled with filtered aCSF. Light pulses were delivered through a light guide microscope illumination system (Lumen Dynamics) modified to accept a blue laser (473 nm, Lasermate Group) in place of the lamp. The laser beam was focused onto the sample through the 60X objective during recordings, with an illumination field of 350μm diameter. Each light pulse was 0.5 ms at 1 Hz, and each recording was an average of approximately ten trials. To minimize the contribution of rundown on the estimation of the plasticity gradient within the striatal slice, recording locations were selected randomly for each slice. For quantification of the ChR2-LFP, each averaged trace was normalized to the peak of the first component. A line was fitted to 10% to 90% of the post-synaptic depolarization phase (red rectangle in Fig 3B) whose slope provides the ChR2-LFP for that recording site. For each slice, the ChR2-LFP slopes across sites were re-scaled from 0 to 1, with the smallest ChR2-LFP set to zero and the largest to 1. For each animal, this normalized mean ChR2-LFP slope for each position along the tonotopic axis was plotted and the plasticity gradient was defined as the slope of the linear fit to this data for that animal. Precaution was taken to select approximately the same slice from every animal to maintain consistency across experiments. To overlay the plasticity gradient maps across animals, the striatal maps for each experiment were aligned to each other using the center of the recording area (as estimated by mean x and y coordinates of recording sites).

### Data Analysis and statistics

All behavior and electrophysiology data were acquired and analyzed using custom designed software written in MATLAB. Wilcoxon Rank Sum Test was performed to test the significance of difference between learning induced plasticity gradients in animals trained on the tonecloud and reversal-tonecloud tasks. Wilcoxon Signed Rank tests were performed to compare training times and performance analysis before and after reversal of animals on the reversal task. Kruskal Wallis test was performed to detect presence of significant differences in the ChR2-LFP slopes along the dorso-ventral axis of the striatum.

